# Migration, proliferation, and elasticity of bladder cancer cells on lectin-coated surfaces

**DOI:** 10.1101/2024.05.12.593008

**Authors:** Marcin Luty, Renata Szydlak, Joanna Pabijan, Ingrid H Øvreeide, Victorien E. Prot, Joanna Zemła, Bjørn T. Stokke, Małgorzata Lekka

**Author notes:** Correspondence; MLekka.

## Abstract

**The alterations in migration, proliferation, and mechanics of cells observed during cancer progression can potentially be linked to enhanced tumor invasiveness. These properties are frequently attributed to the ability to form distant metastasis; however, the direct mutual connection between these properties is not always proven. Here, we studied the migratory, proliferative, and mechanical phenotype of three bladder cancer cells originating from various stages of cancer progression, i.e., non-malignant cell cancer of ureter (HCV29 cells), bladder carcinoma (HT1376 cells) and transitional bladder carcinoma (T24 cells). The results were linked with the organization of actin filaments because of their major role in cell migration. The results classified cells into non-malignant, non-invasive, and invasive, revealing the significant impact of actin filaments in bladder cancer invasion. Based on the reported changes in cancer cell glycosylation, the potential applicability of the observed cancer-related changes to identify invasive cells was demonstrated for the lectin-coated surfaces, which is the potential surface modification for biosensors.**

## 1. Introduction

Cell migration is a fundamental phenomenon accompanying various cellular processes such as immune response [1], morphogenesis [2], or wound healing [3]. Migrating cells use the actin network to probe and move through surrounding environments to guide cells toward specific directions, using two types of actin networks at the leading edge, the lamellipodium and the lamellum [4,5]. The lamellipodium, composed of dynamically changing actin filaments, extends the leading edge by incorporating new monomers close to the plasma membrane. Then, the lamellum, located behind the lamellipodium, forms a more stable structure composed of bundled actin filaments oriented parallel to the leading edge and anchored to the substrate by focal adhesions [6]. Cell migration has detrimental effects when involved in pathological processes such as cancers. Invasion is directly linked with enhanced proliferation, migratory phenotype, and increased deformability of cancer cells. The deregulation of these cellular features during cancer progression is linked with the capacity of tumor cells to escape from the primary tumor site, inducing the formation of metastatic sites at distant body organs [7,8].

Changes in the biomechanical or rheological properties of cells affect biological functions and occur in various abnormalities such as diabetes [9], multiple sclerosis [10], muscular dystrophy [11], or cancer [12]. In most cancers like bladder [12,13], ovarian [14], kidney [15], pancreas [16], breast [17], or melanoma [18], cells soften, while in some cases like colorectal [19] or colon cancers [20], stiffening is observed. The cell mechanics is frequently characterized by an elastic (Young’s) modulus. Its lower value indicates cell softening, while stiffening is shown by a higher modulus than the reference (healthy or non-malignant) cells. Notably, the measurements of the mechanical properties of cells reflect the status of the cell cytoskeleton, particularly the organization of actin filaments [13,21]. Therefore, by accessing cell mechanics, one can get information about the mechanical status of the F-actin network. Softening of cancer cells is frequently related to disorganized actin filament organization, including lack of stress fibers [21–23]. Although most cancer cell types have demonstrated cancer-related softening, only a few studies were related to evaluating the relationship between cell mechanics and invasiveness. Most frequently, the relation between cell mechanics and invasiveness has been related to the stage of the cancer progression but without accompanying experiments showing proliferative or migrative cellular properties [18,24]. Only a few papers showed a direct relationship between the cancer cells and invasiveness for ovarian [25] and pancreatic [16] cancers.

Lectins are plant proteins that identify carbohydrate moieties (i.e., glycans) attached to various cell surface proteins and lipids in a specific way [26]. Therefore, they become essential molecular and cell biology, immunology, pharmacology, medicine, and clinical analysis research tools [27]. In recent years, the use of lectins as a screening tool for potential biomarkers linked with structural changes of glycans in cancers [28]. We have recently shown that specific lectins can be applied to selectively capture and identify label-free bladder cancer cells [29]. These and other studies [30,31] stimulate investigations of how cellular properties such as migration, proliferation, or mechanics change in response to lectin type. Here, we focus on the migratory, proliferative, and elastic properties of bladder cancer cells, emphasizing their behavior on lectin-coated surfaces. These properties were measured for urothelial cell lines (non-malignant HCV29 and carcinoma HT1376 and T24 cells) cultured on the surface coated with previously studied plant lectins from *Dolichos Biflorus*, *Lens Culinaris*, *Phaseolus vulgaris*, and *Triticum vulgaris (wheat germ)*. The choice of lectins was dictated by their affinity to specific glycans present on the surface of bladder cancer cells [32–35]. The glycosylation pattern in cancer (including urothelial cancers) may significantly impact their invasive potential by, e.g., promoting the epithelial-mesenchymal transition [36] or affecting integrins that regulate the metastasis [37]. Glycans, widely expressed in the ECM and on the cell surface, strongly influence cell proliferation, survival, and metastasis during cancer. Therefore, before using lectin-based biosensors coupled to cell mechanics for cancer-related diagnostic or prognostic purposes, the assessment of cell proliferative, migratory, and mechanical phenotype has to be elaborated.

## 2. Materials and Methods

### 2.1 Lectins

Commercially available plant lectins, i.e., *Dolichos Biflorus Agglutinin* (DBA), *Lens Culinaris Agglutinin* (LCA), *Phaseolus vulgaris* (PHA-L), and *Wheat Germ Agglutinin* (WGA) were purchased in Vector Laboratories (Biokom, Janki, Poland) were used. Lectins were dissolved in phosphate-buffered saline (PBS, Sigma-Aldrich, Poznań, Poland) at 125 µg/mL concentration.

### 2.2 Cell lines

Three types of urothelial cell lines were used in this study, namely, non-malignant cell cancer of the ureter (reference cell line HCV29, established in Institute of Experimental Therapy, Wroclaw, Poland), bladder cell carcinoma (HT1376, grade III, ATCC, LGC Standards) and transitional cell carcinoma (T24, ATCC, LGC Standards). The HCV29 and the T24 cells were grown in Roswell Park Memorial Institute Medium 1640 (RPMI 1640, Sigma-Aldrich, Poznań, Poland) supplemented with 10% fetal bovine serum (FBS, ATCC, LGC Standards, USA). The HT1376 cells were cultured Eagle’s Minimum Essential Medium (EMEM, LGC Standards, USA) supplemented with 10% FBS. The cells were cultured in the plastic 25 cm² culture flasks (TPP, Genos, Poland) at 37 °C in a 95% air/5% CO_2_ atmosphere in the CO_2_ incubator (Nuaire, USA). All cell lines were routinely observed under an optical microscope (CK53, Olympus, Poland). The cells were passaged twice a week. Cell line authentication was performed using the FTA Sample Collection Kit for Human Cell Authentication Service (LGC Standards, USA).

### 2.3 Covering the surface with lectins

The surface of 12-well plastic dishes (TPP, Genos, Poland) or 18×18 mm glass coverslips (Bionovo, Legnica, Poland) was coated by incubation the 125 µg/ml solution of DBA, LCA, PHA-L, or WGA lectin in PBS for 2 h. Then, the solution of lectins was removed, and the surfaces of the plastics and slides were dried under a laminar chamber.

### 2.4 Elasticity measurements by atomic force microscope (AFM)

For AFM-based elasticity measurements, cells were seeded onto the surface of 18 × 18 mm glass coverslips placed in 6-well plates (TPP, Genos, Poland) at 40000 cells/well density for 72 hours in the appropriate culture medium (either RPMI 1640 or EMEM). The surface of glass coverslips was coated with lectins. As a control, the bare surface of the glass coverslip was used.

The mechanical properties of bladder cancer cells were measured using AFM (Xe-120, Park Systems, Korea) working in a force spectroscopy mode with commercially available silicon nitride cantilevers with a nominal spring constant of 0.03 N/m (MLCT-D, Bruker, USA). To calibrate the sensitivity of the photodetector, force curves (i.e., dependences between the cantilever deflection and relative scanner position) were acquired on a stiff, non-deformable glass surface. The cantilever spring constants were calibrated using the thermal tune approach [38]. Afterward, force curves were recorded within the nuclear region of the cells. Measurements were repeated 3 times for each condition. The number of cells varied from 14 to 45 cells, depending on the lectin and cell types. For each cell, 25 force curves were acquired in the scan area of 5 µm × 5 µm (a grid 5 × 5 was set).

### 2.5 Young’s modulus determination

Young’s modulus quantified the mechanical properties of living cells, describing the resistivity of cells to compression. In terms of cells, its value is used to quantify the ability of cells to deform, i.e., their deformability [39]. As already described elsewhere [40], force curves were converted to force versus indentation curves and fitted with Hertz-Sneddon contact mechanics for conical tip shape [41] using the following equation:

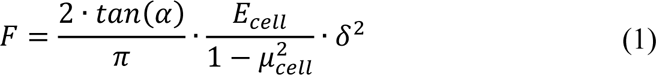

Where *F* is the load force, δ is the indentation depth, *µ_cell_* is the Poisson’s ratio (assumed to be 0.5 for incompressible materials), α the half angle of the indenter tip, and *E_cell_* is Young’s modulus of the cell. The final Young’s modulus, calculated from the indentation depth up to 500 nm, was expressed as a mean ± standard deviation (SD).

### 2.7 Proliferation assay

Cells were seeded in 24-well plates (TPP, Genos, Poland) at the density of 4000 cells/mL of the corresponding culture medium supplemented with 10% FBS (0.5 mL of cell suspension per well). Cells were grown for a maximum of 120 h. At each time point (i.e., 24 h, 48 h, 72 h, 96 h, or 120 h), the cells (from 3 wells) were passaged and counted using a Bürker chamber (Marienfeld, Germany). The proliferation rate of bladder cancer cells was quantitatively assessed by a population doubling time (PDT) by fitting the following equation to the obtained data:

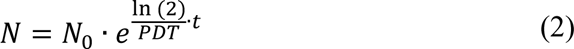

where *t* is the culture time (days), *N* is the final number of cells, and *N_0_* is the number of cells on day 0. The experiments were repeated 3 times for each time. In one experimental run, the number of cells was counted from 3 wells. The determined means ± standard deviations were plotted as a function of culture time. The final PDT value was obtained from the fit of equation (2); thus, error bars are a standard error.

### 2.8 MTS assay

The MTS colorimetric test (Promega, Poland) was used to determine the influence of lectin-coated substrates on the proliferation of bladder cancer cells. HCV29, T24, and HT1376 cells were seeded on 24-well plates (TPP, Genos, Poland) at 4000 cells/mL density in 0.5 ml of corresponding culture medium supplemented with 10% FBS. Next, cells were cultured at 37 °C in the 5% CO_2_/95% air atmosphere in the CO_2_ incubator (Nuaire, USA). At each time, i.e., 24 h, 72 h, or 120 h, the 0.5 ml of the culture medium was exchanged into a fresh one, and 50 µl of tetrazole salt was added. Then, cells were incubated for the next 2 h at 37 °C. A yellow-colored tetrazole salt is taken up by living cells by pinocytosis. As a result of the oxidoreductive activity of mitochondria, the tetrazole salt is reduced by living cells to colored, purple formazan products, which are fully soluble in culture media. The amount of dye can be assessed spectrophotometrically. Therefore, after 2 h of incubation, the absorbance at a wavelength l = 490 nm was measured using an LEDETECT 96 (Labexim Products, Austria) microplate reader. The MTS assay was repeated three times for each time point and cell line. The final PDT value was obtained from the fit of equation (2) to the relation between ELISA readout and time (error bars are a standard error).

### 2.9 Single-cell migration assay

Cells were seeded onto the surface of 12-well plates (TPP, Genos, Poland) with a plain or lectin-coated surface. Cells (40000 cells per well) were cultured in the corresponding medium (RPMI 1640 for HCV29 and T24 cells or EMEM for HT1376 cells) for 24 hours. A set of images of migrating cells was taken, at regular intervals of 10 minutes, for 4 hours using videoscopy mode provided by the IX83 inverted Olympus microscope (Olympus) every 10 minutes apart over 4 hours. The obtained images were analyzed using the Hiro program (courtesy of the Department of Cell Biology at the Faculty of Biochemistry, Biophysics, and Biotechnology of the Jagiellonian University, Cracow, Poland). Motion trajectories are presented as circular diagrams in which the starting points of the trajectories are reduced to a common origin located at the origin of the coordinate system [42]. The mean single-cell velocity was calculated to quantify the cell migration activity. Then, statistical analysis was performed from 3 independent replicates, in which the trajectories of 150 cells were analyzed.

### 2.10 Wound healing assay

Scratch-making inserts (Culture-Insert 2 Well, IBIDI) were placed on the surface of 12-well plates. Cell suspension (50,000 cells per insert compartment in 70 µl of cell medium supplemented with 1% FBS). After 24 hours, the inserts were removed, and 1 ml of the culture medium supplemented with 1% FBS was added to initiate the assay (t = 0 hours). Optical images were subsequently recorded at 1, 3, 6, 12, and 24 hours. The wound healing rate was calculated from the mean distance between cell monolayer borders measured using Image J software (version 1.53k, https://imagej.nih.gov/ij/, 6 July 2021). The experiments were repeated 3 times. In one experimental run, 10 to 20 randomly selected locations were evaluated, for which a mean was calculated. The final value of the wound closure was expressed as a mean ± standard deviation from *n* = 3-4 replicates.

### 2.11 Transmigration of single cells through matrigel

The surface of the Corning Transwells inserts (Sigma-Aldrich, Poland, Poznań) with pore sizes of 8 µm was coated in two ways. First, a solution of ECM gel from Engelbreth-Holm-Swarm murine sarcoma (300 µg/mL, Sigma-Aldrich, Poznań, Poland; referred to here as matrigel) was placed (100ul ECM solution per transwell) and incubated at 37°C for 3 hours. The solution was removed, and the inserts were dried in a laminar flow chamber (Nuaire, USA). Human clabber cells (HCV29, HT1376, and T24 cells) were harvested, centrifuged, and suspended in 5 µM Cell Tracker Red CMPTX dye (Invitrogen, Thermo Fisher Scientific, Waltham, MA, USA), then incubated at 37°C for 60 minutes, after which cells were washed with PBS and centrifuged. The cell pellet was suspended in the corresponding culture medium supplemented with 1% FBS (5·10^4^ cells/mL). Such prepared cell suspensions (100 µl) were transferred to the inserts and placed in the wells containing 600 µl of culture medium supplemented with 10% FBS, followed by 24-hour incubation in the CO_2_ incubator. The BSA-containing (50 mg/mL) culture medium was used as a negative control. After this time, the bottom surface of the whole well was scanned and imaged using a fluorescence microscope (IX83, Olympus). The number of cells that migrated through the matrigel was determined as mean ± standard deviation derived from *n* = 3 repetitions.

### 2.12 Visualization of actin filaments in single bladder cancer cells

Two main cytoskeletal components were stained fluorescently, i.e., F-actin and microtubules. The cells were cultured in 12-well plates in the corresponding culture medium supplemented with 10% FBS for 48 hours. Next, the cells were fixed by adding a 3.7% paraformaldehyde solution in PBS for 10 min and washed twice with the PBS buffer. Next, the cell membrane was permeabilized by incubating the cells with a cold 0.2% Triton X-100 (Sigma) in PBS solution for 5 min. Finally, cells were rinsed with PBS buffer. Such prepared coverslips with cells were fluorescently labeled. First, microtubules were stained with Alexa Fluor 555 labeled anti-tubulin antibody (Molecular Probes, Thermo Fisher Scientific, Waltham, MA, USA) dissolved in PBS buffer (1:200) for 24 hours. Then, the sample was washed with PBS buffer, and F-actin was labeled with phalloidin conjugated Alexa Fluor 488 (Molecular Probes, Thermo Fisher Scientific, Waltham, MA, USA) dissolved in PBS buffer (1:200) for 60 min, followed by rinsing it with PBS buffer. The cell nuclei were stained with Hoechst 33342 dye (Sigma-Aldrich, Poznań, Poland) dissolved in PBS buffer (1:5) for 15 min. Samples were visualized with a confocal microscope (Zeiss Axio Observer.Z1/7 with a detector LSM80 GaAsP-PMT). Two channels were recorded: green (Alexa Fluor 488; excitation 493 nm and emission 517 nm) and blue (DAPI; excitation 355 nm and emission 465 nm).

### 2.13. Fluorescent visualization of actin filaments in cells cultured on the lectin-coated surfaces

Immunostaining of F actin was conducted as follows. The cells were grown on 12-well plates with an RPMI-1640 or EMEM medium supplemented with 10% FBS for 48 hours. Next, the cells were fixed by adding a 3.7% paraformaldehyde solution dissolved in the PBS buffer for 10 min and then washed twice in the PBS. The cell membrane was permeabilized by incubating the cells with a 0.2% Triton X-100 (Sigma) in PBS solution for 5 min. Finally, cells were rinsed with PBS buffer. Such prepared coverslips with cells were fluorescently labeled. The actin filaments were stained with phalloidin fluorescently labeled with Alexa Fluor 488 (Invitrogen) dissolved in PBS buffer (1:200) for 60 min. Then, the sample was washed with PBS buffer. Samples were visualized with an inverted optical microscope (Olympus IX83) with a fluorescence module. A 100W mercury lamp was applied to excite Alexa Fluor 488 dye. Images were acquired using the ORCA-spark (Hamamatsu Photonics, Hamamatsu, Japan) digital camera, providing a 2.3 megapixel 1920 px × 1200 px image.

### 2.13 Statistic

If not stated explicitly, the statistical significance was obtained from the unpaired t-test at the significance level of 0.05. The F-test was used to determine whether the two PDT datasets fitted with the same model differed significantly. Notation: *ns* – not statistically significantly different, *\*p* < 0.05, *\*\*p* < 0.01, *\*\*\*p* < 0.001.

## 3. Results

### 3.1. Intrinsic mechanical and migratory properties of bladder cancer cells

In our study, we focused on urothelial cell lines originating from three stages of cancer progression, namely, non-malignant HCV29 cells that originate from the ureter [43,44] and two carcinoma cell lines (HT1376 and T24 cells) with different cancer grades. The former is an epithelial-like cell isolated from the bladder of a white, 58-year-old female classified as carcinoma with grade 3 [45], while the latter is a similar epithelial-like cell line isolated from the bladder of a white, 81-year-old female classified as transitional cell carcinoma with grade 3/4 [46]. In the first steps, these cells were cultured on glass coverslips or tissue surface plastic (Petri dish) correspondingly for AFM and fluorescent microscopy.

The deformability of cells was obtained using AFM-based elasticity measurements. The results showed that non-malignant HCV29 cells are stiffer than the cancerous cells (**Fig. 1a**, **Table 1**; *p* = 0.0007 (HCV29/HT1376) and *p* = 0.0471 (HCV29/T24)). Among the cancer cell lines, HT1376 cells were the softest ones. These results agreed with our previous results [12,13,22,47].

**Figure 1.**
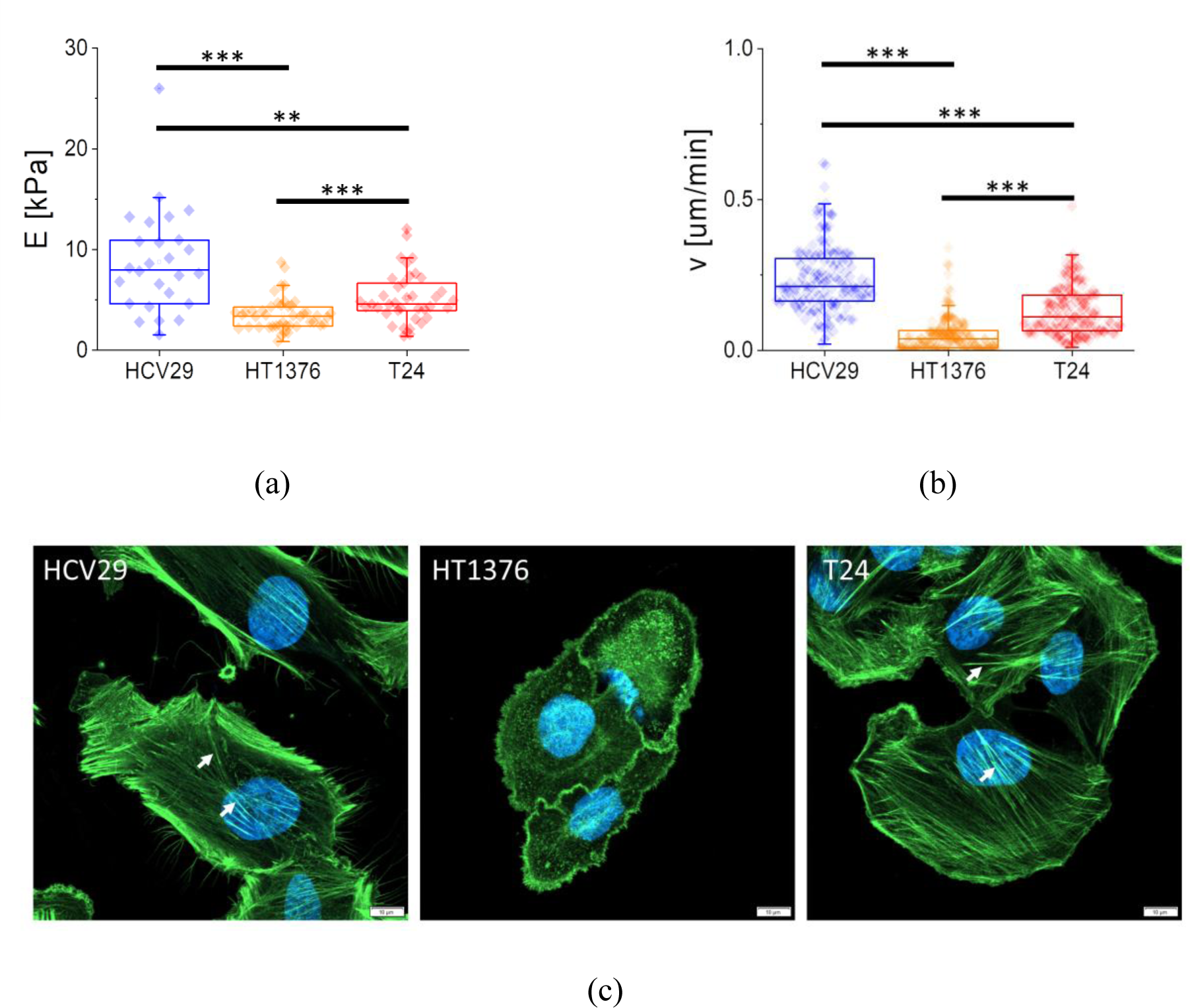
Mechanical, migratory, and structural aspects of bladder cancer cells. (**a**) Deformability of the cells expressed as Young’s modulus value in kPa. Each point denotes the mean value of Young’s modulus determined for a single cell (total number of cells n = 26, 45, 38 cells for HCV29, HT1376, and T24 cells, respectively). (**b**) Migration speed in micrometers per minute. Each point denotes a speed value obtained for the individual cell (n = 150 cells for each cell type in 3 replicates). (**c**) Fluorescent images of F-actin organization were visualized using phalloidin conjugated with Alexa-Fluor 488 fluorophore, while cell nuclei were stained with Hoechst 33342 (white arrows indicate stress fibers). The scale bar is 10 µm.

**Table 1.** Mechanical properties of human urothelial cells.

| Cell line | Young's modulus<br>(mean ± SD (number of cells)) |
| --- | --- |
| non-malignant HCV29 cells | $8.8 \pm 1.7$ kPa (26 cells) |
| carcinoma HT1376 cells | $3.6 \pm 0.5$ kPa (45 cells) |
| transitional cell carcinoma T24 cells | $5.2 \pm 0.9$ kPa (38 cells) |

In parallel, the migratory properties of these cells were assessed using an inverted optical microscope to monitor changes in the migration velocity of the cells (**Fig. 1b**, cells displacements are shown in **Suppl. Fig. S1**). The determined speed of migration was the following: 0.24 ± 0.11 µm/min (*n* = 150 cells, HCV29 cells), 0.05 ± 0.06 µm/min (*n* = 150, HT1376 cells), and 0.13 ± 0.08 µm/min (*n* = 150, T24 cells). The lowest migration speed was observed for HT1376 cells, while the largest was for HCV29 cells.

Knowing that the elasticity reflects the status of the cell cytoskeleton [21,48], F-actin was fluorescently labeled with phalloidin conjugated with Alexa Fluor 488 dye (emission 493 nm/excitation 518 nm), and its intracellular organization was recorded for all three types of cells (**Fig. 1c**). Phalloidin binds to the polymerized form of actin, not to actin monomers. Fluorescent images reveal thick F-actin bundles, probably stress fibers, inside non-malignant HCV29 and cancerous T24 cells but not inside cancerous HT136 cells. Mostly, short filaments (detectable as a shadow in fluorescent images) were formed in these cells.

The results show that the deformability of cancer cells and migration of bladder cancer cells were related to the organization of F-actin filaments. The deformability of cells is intuitively related to the cell cytoskeleton, while such a relation for the link between cell migration and the actin cytoskeleton is not always clear. The migration of cells is strongly linked with the actin filaments [49,50]. Cancerous HT1376 cells displaying a disorganized cytoskeleton and lack of stress fibers were the softest cells with poor migration ability. Well-developed actin cytoskeleton in non-malignant HCV29 cells manifested in the largest rigidity and highest migration ability

#### 3.1.1. Invasiveness of bladder cancer cells

Enhanced cell migration of cancer cells is associated with cancer invasiveness [51,52]. Expecting various behaviors of bladder cancer cells on lectin-coated surfaces that could be linked with the invasiveness, in our next step, we assessed the ability of bladder cells to invade by various methods, such as proliferation rate, wound healing, transmigration of single cells through matrigel and monolayer of HCV29 cells (**Fig. 2**), and transmigration of cancer spheroids through HCV29 monolayer (**Suppl. Note S1, Suppl. Fig. S2**).

**Figure 2.**
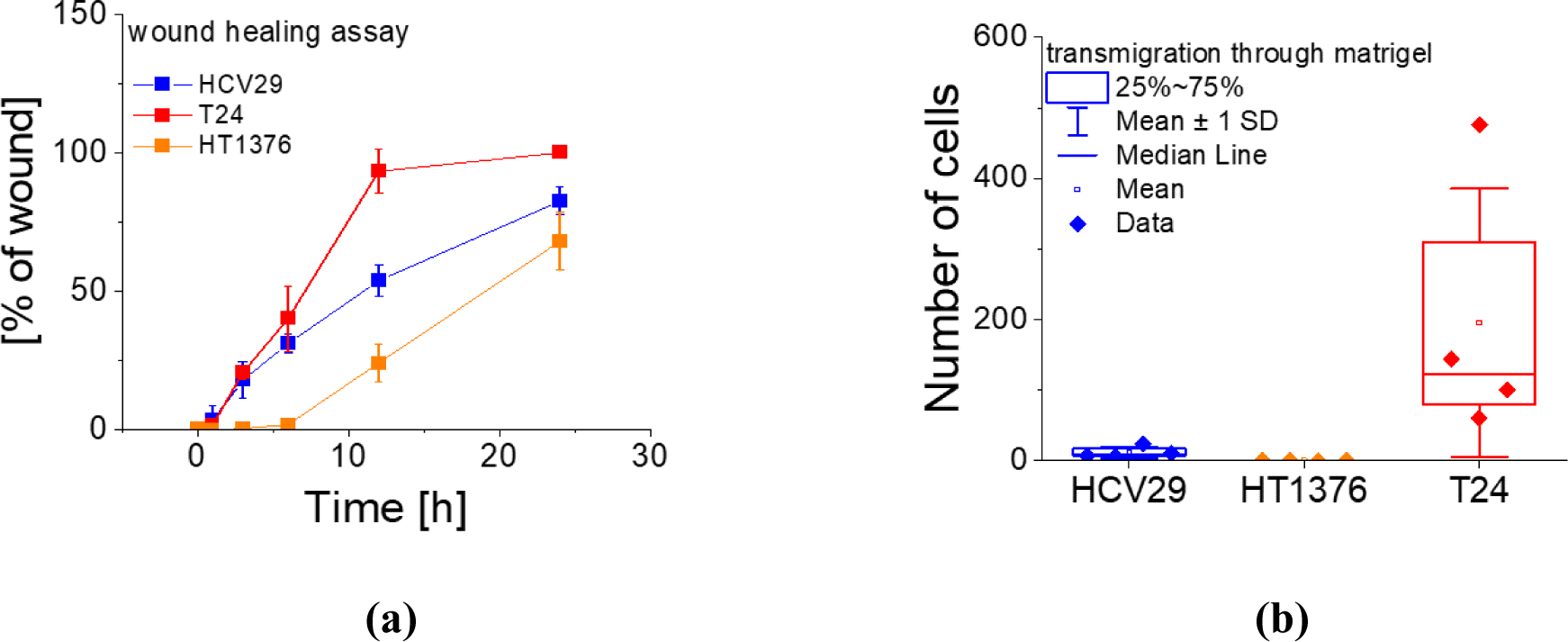
Transmigration assessment to characterize invasiveness of bladder cancer cells: (**a**) wound healing assay quantifying the invasion of cells on a 2D surface. Each point denotes a mean ± s.d. calculated from n = 40 to 60 locations. (**b**) Transmigration of bladder cancer cells through the matrigel mimicking the ECM environment. Each data point represents the total number of cells that migrate through one transwell, n = 4 repetitions).

The proliferation rate can be described by PDT, after which the number of cells doubles. PDT for the studied cell line was 24.2 ± 0.6 h (coefficient of determination, COD = 0.99859), 30.8 ± 0.5 h (COD = 0.9995), and 17.0 ± 0.1 h (COD = 0.99984) for non-malignant HCV29, cancerous HT1376, and T24 cells, respectively. The proliferation rate correlates with the results obtained from the wound healing assay applied here (**Fig. 2a**). It describes the ability of the cells to migrate on the 2D surface (the bottom surface of the Petri dish). It is quantified by the surface area occupied by cells during culture time (the percentage of wound closure). The results show that the fastest wound closure was observed for cancerous T24 cells and the slowest for HT1376 cells.

Cell transmigration assay was conducted to evaluate the invasiveness of the studied bladder cancer cells. First, the ability of all three types of cells to migrate through the matrigel layer was evaluated after 24h (**Fig. 2b**). Matrigel mimics the ECM surrounding as it is a well-defined soluble basement membrane extract [53]. After 24 hours, on average, 11.5 ± 7.9 non-malignant HCV29 cells and 195 ± 190 cancerous T24 cells passed to the bottom surface. No single cells were observed for cancerous HT1376 cells. Further, we evaluate the capability of cancer cells (HT1376 and T24) to migrate through a monolayer of non-malignant HCV29 cells (**Suppl. Fig. S2a**). The relation was similar, i.e., HT1376 did not pass, while T24 cells passed through it. However, in the latter case, the number of cells was much smaller. A drop from ∼200 to 43 cells was observed (mean equaled 42.7 ± 2.5 cells).

The data indicate that the T24 is the cell type with the highest migratory phenotype, which also results from the strong interplay with the proliferation rate. Fluorescent images of the cancer spheroids, prepared from HT1376 and T24 cells, confirmed that HT1376 cells could not transmigrate through the HCV29 cell monolayer (**Suppl. Fig. S2b**). One can observe cells escaping from cancer spheroid formed from T24 cells but not from spheroids formed from HT1376 cells. The results show that cancerous T24 cells were characterized by the highest motile phenotype and ability to transmigrate through matrigel and the monolayer of non-malignant HCV29 cells among the studied cancer cell lines. Such behavior allows us to conclude that these cells are invasive while HT1376 cells are not.

### 3.2. Proliferation of bladder cells cultured on lectin-coated surfaces

The MTS assay was conducted to elaborate on how lectins used for surface coating affect the proliferation activity of bladder cells (**Suppl. Fig. S3**). For each culture duration, the mean absorbance values and standard deviation were calculated from 9 readouts (3 repetitions; in a single run, three wells were measured), followed by plotting them as a function of culture time. Such data were fitted with eq. (2) to obtain the PDT value for each pair composed of the specific cell and lectin type (**Suppl. Fig. S3a** presents an exemplary fitted curve). The obtained PDT for bladder cancer cells cultured on lectin-coated surfaces is presented in **Table 2**.

**Table 2.** Population doubling time was determined using MTS assay for bladder cancer cells cultured on a lectin-coated surface (expressed in hours), resulting from the fit of equation (2); Supplementary Figure S3. The number of cells was assessed from n = 9 readouts.

| Cells | Control | DBA | LCA | PHA-L | WGA |
| --- | --- | --- | --- | --- | --- |
| non-malignant HCV29 | $26.1 \pm 2.2$ h | $25.7 \pm 4.2$ h | $26.8 \pm 3.1$ h | $24.6 \pm 3.5$ h | $29.3 \pm 2.6$ h |
| invasive T24 | $17.0 \pm 4.5$ h | $15.3 \pm 1.2$ h | $15.1 \pm 1.2$ h | $13.7 \pm 1.8$ h | $11.0 \pm 5.3$ h |
| non-invasive HT1376 | $36.1 \pm 3.7$ h | $32.5 \pm 13.9$ h | $31.4 \pm 8.8$ h | $30.3 \pm 0.6$ h | $29.4 \pm 17.9$ h |

No statistical significance was detected when applying the F-test to compare datasets fitted with one model (at 0.05 level). Despite that, the PDT determined for non-malignant HCV29 cells revealed the trend to be insensitive to DBA and LCA coatings (PDT changes about 1-3% in relation to control cells) and showed faster proliferation when cultured on the PHA-L-coated surface (PDT dropped by about 6%). Their growth on the WGA-coated surface resulted in slower proliferation (PDT increased by ∼3 hours). Statistical significance (at the level of 0.05) was obtained for specific lectin coatings and cancer cell types (**Suppl. Fig. S3**). The PDT for non-invasive HT1376 cells was longer than for HCV29 cells, i.e., it changed from 29 to 36 hours. The longest PDT was observed for control conditions (HT1376 cells on the Petri dish surface, 36.1 ± 3.7 h). It is reduced for these cells cultured on lectin-coated surfaces, subsequently as *PDT_DBA_ > PDT_LCA_ > PDT_PHA-L_ > PDT_WGA_* (*p <* 0.001 in relation to control for all lectins). Such a relation indicates that lectins fasten cell division, showing that they can be used to stimulate proliferation. Such an effect has already been reported for lymphocytes [54]. Among lectin coatings, a significant difference in PDT for non-invasive HT1376 was observed for two pairs: *PDT_LCA_* (26.8 ± 3.1 h) and *PDT_PHA-L_* (24.6 ± 3.5 h) with *\*\*p* < 0.01 and *PDT_PHA-L_* (24.6 ± 3.5 h) and *PDT_WGA_* (29.3 ± 2.6 h) with *\*\*\*p* < 0.001. The lowest proliferation time was obtained for the PHA-L coated surface. The invasive T24 cells were characterized by the smallest PDT compared to the other two cell lines in control conditions. Similar to non-invasive HT1376 cells, the PDT for T24 cells decreases with applied lectin coatings, indicating analogous faster cell division. The decrease follows a similar relation, i.e., *PDT_DBA_ ≅ PDT_LCA_ > PDT_PHA-L_ > PDT_PHA-L_* (in relation to control cells). The *PDT*_WGA_ value was burdened by a large standard error, resulting in the lack of statistical significance compared to PDT values for cells coated with other lectins but not with control. Among lectin coatings, a significant difference in PDT for invasive T24 was observed between the following pairs: *PDT_DBA_* (15.3 ± 1.2 h) and *PDT_LCA_* (15.1 ± 1.2 h) with *\*p* < 0.05, *PDT_DBA_* (15.3 ± 1.2 h) and *PDT_PHA-L_* (13.7 ± 1.8 h) with *\*\*\*p* < 0.001, and *PDT_LCA_* (15.1 ± 1.2 h) and *PDT_PHA-L_* (13.7 ± 1.8 h) with *\*\*\*p* < 0.001. Altogether, these results show that the lowest proliferation time was obtained for the PHA-L coated surface, regardless of the cell type studied. The WGA coating resulted in the large PDT variability observed in cancer cells, making this lectin unfavorable for the surface coating.

### 3.3. Migration of bladder cancer cells on lectin-coated surfaces

Next, we wanted to answer whether migratory properties are related to the proliferation rate. The migration of bladder cancer cells was obtained analogously to that of control cells (i.e., grown on a Petri dish surface). First, we compared the final distributions of the migration speed for control cells (**Fig. 3a**).

**Figure 3.**
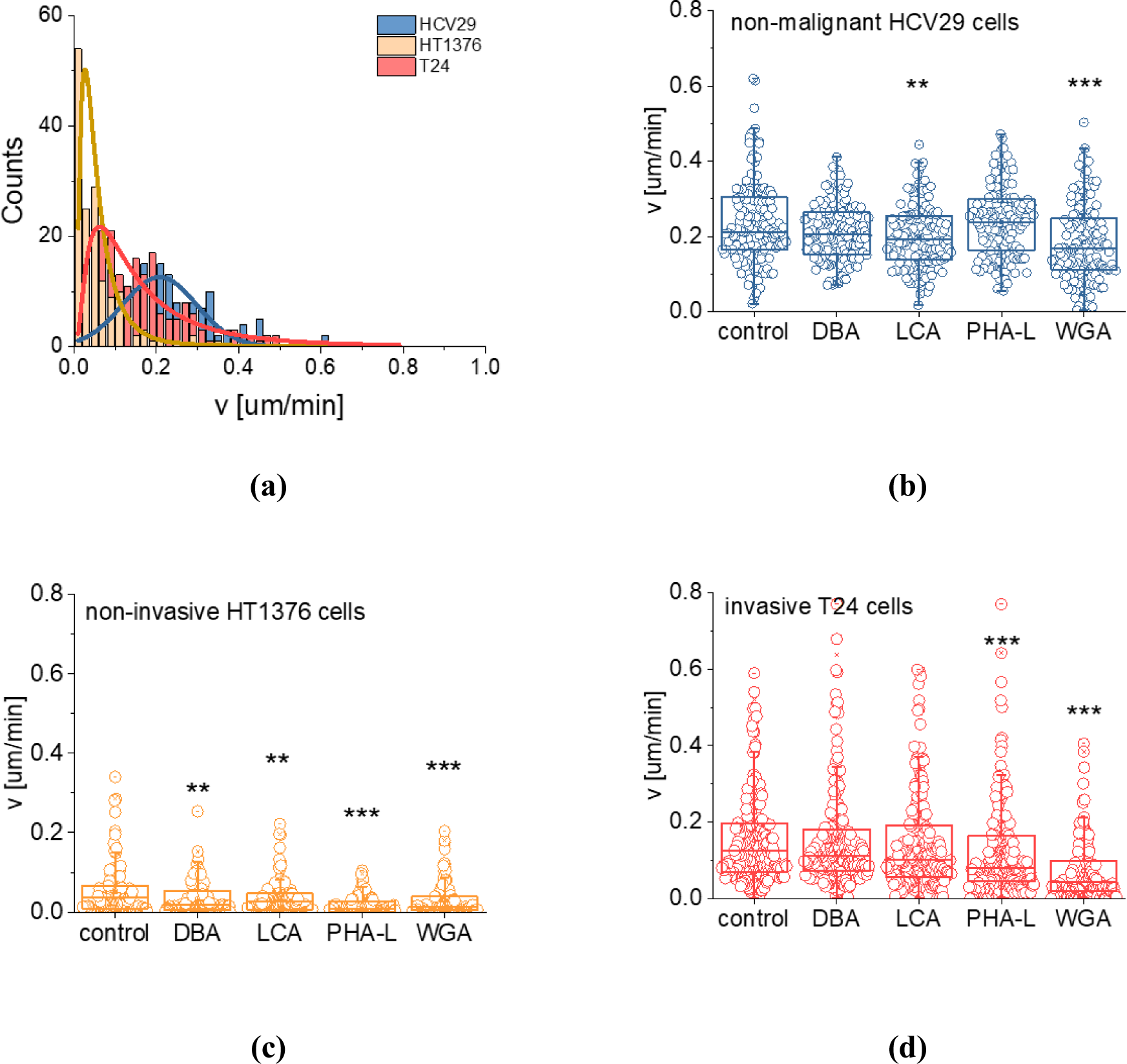
Migration speed for bladder cancer cells on lectin-coated (DBA, LCA, PHA-L, and WGA) surfaces. (a) Exemplary distribution of migration speed for all studied cell lines (control, Petri dish surface, no lectins involved) with fitted lognormal function. Box plots of migration speed for (b) non-malignant HCV29, (c) non-invasive HT1376, and (d) invasive T24 cells. Statistical significance relative to the control surface for each cell line was evaluated using the non-parametric Mann-Whitney test at the significance level of 0.05 (notation: *p < 0.05, **p < 0.01, ***p < 0.001). On average, 150 cells were analyzed for each cell and lectin type.

The distributions were characterized by lognormal function, indicating that among each cell population, a subpopulation of cells migrates faster than most of the cells.

The migration speed for non-malignant HCV29 cells ranged from 0.18 to 0.24 µm/min, depending on the lectin type used for surface coating (**Suppl. Table S1**). Statistical significance was obtained for cells with lower migration speed, i.e., for HCV29 cells migrating on LCA or WGA-coated surfaces: 0.20 ± 0.08 µm/min (*p* = 0.00965) and 0.18 ± 0.10 µm/min (p < 0.001) compared to control cells migrating on Petri dish surface (0.24 ± 0.11 µm/min). Non-invasive HT1376 cells were sensitive to the presence of each lectin type. Their migration speed dropped from 0.05 ± 0.06 µm/min (migration of cells on uncoated Petri dish surface) to 0.02-0.03 µm/min for each lectin type. Invasive T24 cells were less sensitive to the lectin presence. Slower migration of these cells was observed for PHA-L and WGA-coated surfaces (0.12 ± 0.12 µm/min and 0.07 ± 0.08 µm/min, respectively). The migratory response of bladder cancer cells (non-malignant HCV29 and cancerous HT1376 and T24 cells) was lectin-type dependent. Notably, slower migration was observed on the WGA-coated surface for all bladder cancer cells. Surprisingly, this observed drop in migration speed was not related to the adhesion of these cells to the WGA-coated surface [29].

### 3.4. Elastic properties of bladder cells on lectin-coated surfaces

The primary receptors responsible for cell attachment to any surface are integrins. These transmembrane proteins are glycosylated [37,55]. As glycans can affect cell adhesion, which is related to the organization of the actin cytoskeleton, we expect that lectins may affect cell mechanics by binding to surface glycans attached to integrins. Therefore, the elastic modulus was measured by AFM for cells cultured on lectin-coated surfaces (**Fig. 4a, Suppl. Table S2**).

**Figure 4.**
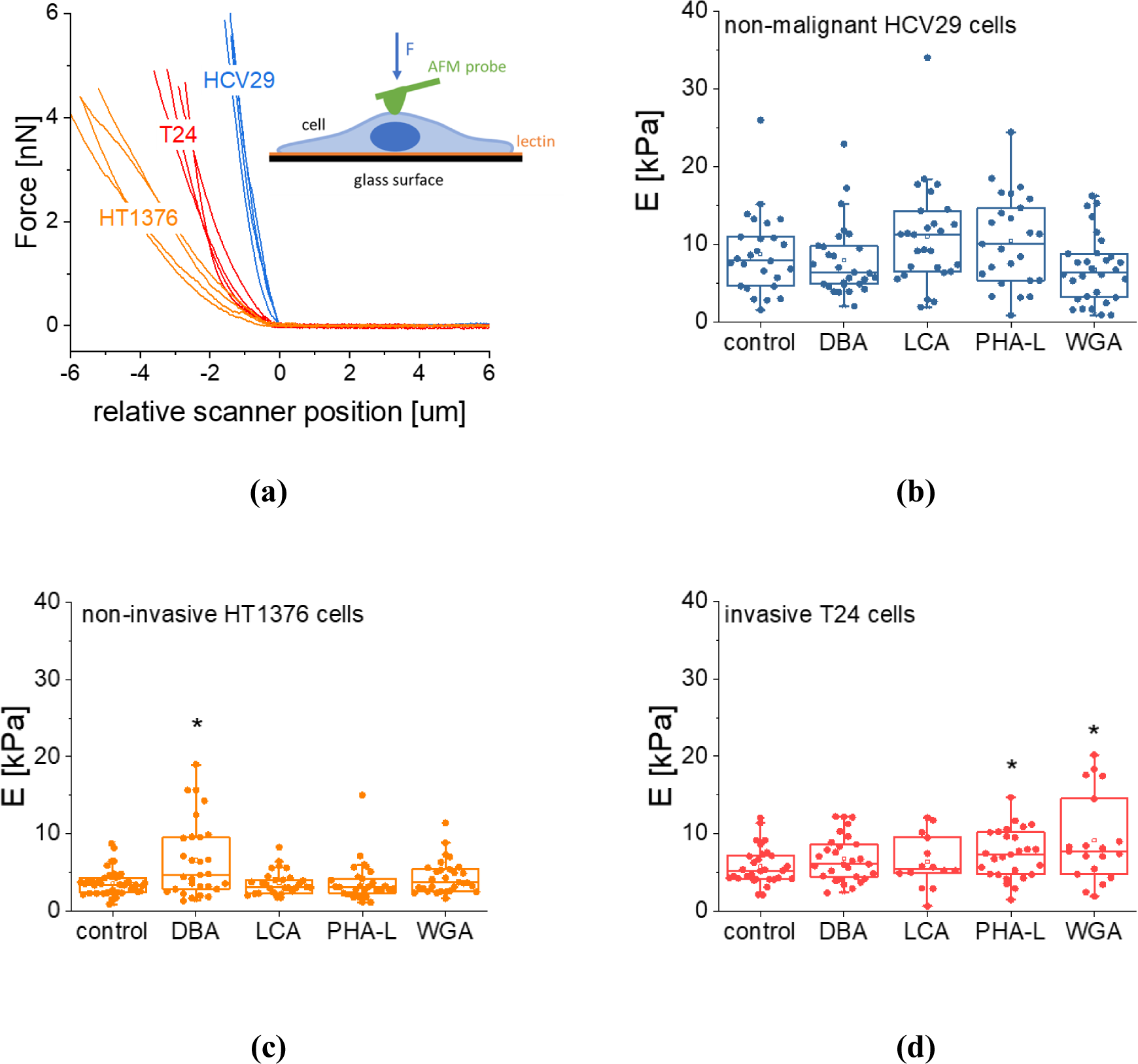
Mechanical properties of bladder cancer cells. (**a**) Exemplary force curves recorded on bladder cancer cells (control). Box plots of Young’s modulus for (b) non-malignant HCV29, (c) non-invasive HT1376, and (d) invasive T24 cells. Statistical significance was evaluated using a non-parametric Mann-Whitney test at the significance level of 0.05 (notation: *p < 0.05)

The presence of lectins on the surface barely affected the elastic properties of the studied bladder cancer cells. The Young’s modulus for non-malignant HCV29 cells showed large variability ranging from 3 kPa to even 35 kPa. Such a large variability does not allow for obtaining statistical significance among the studied cells cultured on various lectins. Notably, the mechanical variability of HCV29 cells was similar, regardless of the lectin used as a surface coating. In the case of non-invasive HT1376 cells, the statistically significant moduli difference was obtained only for the DBA-coated surface (*p* = 0.0117), although a substantial modulus variability was observed. The altered mechanical properties for invasive T24 cells were observed for PHA-L (*p* = 0.03779) and WGA (*p =* 0.02943) coated surfaces.

### 3.5. Actin cytoskeleton organization in bladder cancer cells cultured on lectin-coated surfaces

In the final step of our study, we visualized fluorescent labeled F-actin to evaluate whether the presence of lectin on the surface used for cell growth affects the organization of actin filaments (**Fig. 5**). The organization of actin cytoskeleton follows the structure of single cells; however, images did not show substantial changes in their organization and distributions inside the cells compared to control cells cultured on uncoated Petri dish plastic surface.

**Figure 5.**
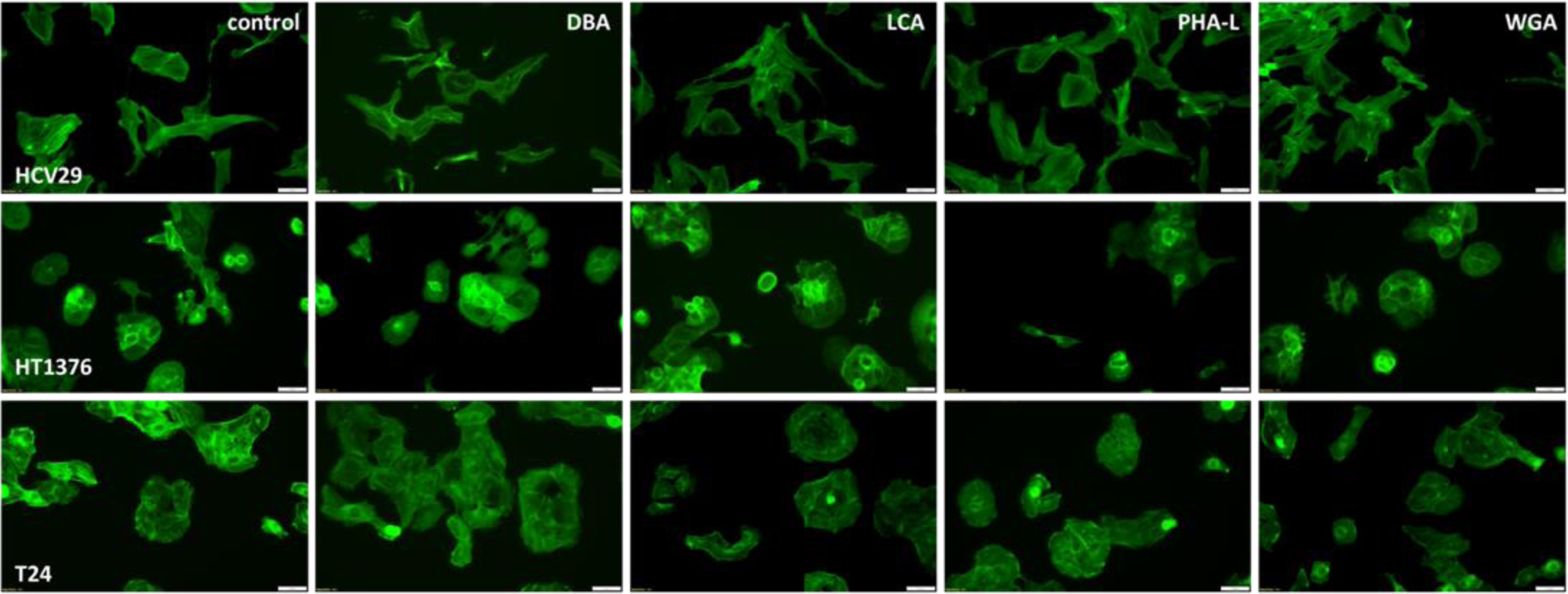
Actin filaments organization of bladder cancer cells. The organization of actin filaments (F-actin stained with phalloidin conjugated with Alexa Fluor 488 dye) inside the bladder cancer cells (HCV29, HT1376, and T24 cells) cultured on uncoated and lectin-coated Petri dish plastic surfaces. The scale bar is 50 µm.

The studied bladder cancer cells revealed no visual changes in the organization of F-actin in response to the lectin coatings, which is considered to be in line with modest changes in the mechanical properties of the studied cells (**Fig. 4**).

## 4. Discussion

The use of biosensors in clinical practice attracts attention due to practical and cost reasons to detect biomarkers that would support the realization of efficient and timely screening [56]. The outcome of such tests enables the precise diagnosis in the early phase of the disease [57], or it could used to sense the effectiveness of treatment [58]. Among many applications, the design of biosensors focuses on their application to detect various cancer-associated biomolecules produced and secreted by tumor cells [59]. Lectins are molecules that possess the attractive potential to be used for label-free identification of cancer due to their specificity to glycans present on the surface of cells as aberrant glycosylation is observed in the disease progression [60–62]. We have recently shown that specific lectins can be applied to selectively capture and identify label-free bladder cancer cells [29]. However, the adhesion of cells to lectin-coated surfaces is only one aspect. Here, we studied how bladder cancer cells behave on lectin-coated surfaces regarding their migratory, proliferative, and mechanical properties.

Cell migration is an essential biological process that affects all aspects of the organism’s proper functioning, from forming distinct tissues to wound healing or targeting various malfunctioning cells or pathogens by immune cells [63]. Unfortunately, the orderly regulated cell migration becomes disturbed in pathological conditions such as inflammatory diseases or cancers [64]. Our results for bladder cancer cells showed the largest migration speed for non-malignant cell cancer of ureter HCV29 cells than for bladder carcinoma T24 cells. The lowest migration speed was observed for bladder carcinoma HT1376 cells. Such relation was observed in the mechanical properties of these cells, i.e., the most rigid were HCV29 cells, then T24 cells, and the softest were HT1376 cells. Thus, rigid bladder cells possess the largest ability to migrate; simultaneously, the softest cells migrate slowly. The actin cytoskeleton organization inside the cells supports these findings. Cell migration and mechanics are governed by the cytoskeleton, a dynamic network of filaments distributed inside the cell that maintains the migratory properties through continuous polymerization and depolymerization of actin filaments in response to inter- and intracellular requests [65,66]. Differences in the arrangements of the actin cytoskeleton have been reported for normal and cancerous cells across a range of cancers, such as breast [67], bladder [22], or thyroid [23] cancers. The ability of the studied cells through a barrier was assessed to verify whether these properties are related to the invasiveness of bladder cancer cells. Unlike expected, the transmigration of T24 cells through matrigel was larger than for HCV29 cells, which indicates that cell migratory properties on 2D surfaces cannot be directly translated to the 3D environment. However, the relation between cancer cells, i.e., low migratory HT1376 and high migratory T24 cells, was preserved. T24 transmigrates through both matrigel and a cellular monolayer composed of HCV29 cells. Intuitively, the reason for that could be the smaller adhesion of T24 cells to HCV29 cells. We could also speculate that actin bundles (probably stress fibers) in T24 cells facilitate their passage by membrane deformation linked with actin polymerization-dependent force production [66].

The migration of bladder cancer cells is strongly linked with their elastic properties and can be related to the organization of actin filaments [49,50]. These properties are not related to the ability of cells to migrate through the matrigel surface and the monolayer of HCV29 cells. Thus, we can conclude that for bladder cancer cells, the largest cell deformability cannot be linked with the higher invasiveness of this cancer type. Our results oppose the belief that the softest can migrate easily and generate the question of why the softest cells with poorly differentiated cytoskeleton are not invasive. In contrast, migrative and stiffer cancer cells displaying thick actin bundles in the cytoskeletal structure are highly invasive. The migration of cancer cells is linked with the actin filaments forming stress fibers [49,50]. These structures are used by cells to form a leading edge. Actin stress fibers are needed to mediate myosin II-based contractility in close cooperation with associated focal adhesion [68]. We speculate that stress fibers are used by cancerous cells to form connections with surrounding cells or ECM. The generated tension helps cells migrate through the densely packed normal cells or ECM layer. Without such additional external forces, the softest cells cannot migrate; thus, stress fibers seem beneficial for cancer cell invasion and dissemination.

Most cell surface receptors like integrins and cadherins are glycosylated, influencing cancer metastasis [69]. They impact cellular signaling and regulate tumor cell-cell adhesion and cell-matrix interactions [69,70]. To elaborate on the effect of glycans on bladder cancer cells, we study the proliferative, migratory, and mechanics of these cells. By adhering cells to lectin-coated surfaces, we influence their integrin-based interactions. Thus, we postulate that by choosing a specific integrin type, we can deduce which glycans participate vigorously in cell-ECM adhesion. The results showed the lectin-dependent relations; however, the strongest effect was observed for the WGA-coated surfaces. For cancer cells, we observed the highest increase in proliferation rate with simultaneous inhibition of migratory phenotype and the largest stiffening (of about 60%) for invasive T24 cells. The WGA influence was slightly smaller but still pronounced for non-invasive HT1376 cells and the smallest for non-malignant HCV29 cells. The lectin WGA (wheat germ agglutinin from *Triticum Vulgaris*) interacts with the sialylated glycoproteins but may also bind to N-Acetyl-D-glucosamine (GlcNAc) [71]. Sialylated glycoproteins have already been reported to be a potential biomarker as they may drive cancer progression, as shown for prostate [72], breast [73], or ovarian [74] cancers.

## 5. Conclusions

The relationship between the organization of the cell cytoskeleton, particularly actin filaments and the migratory and mechanics of cells, is crucial to understanding how cancer cells behave and become more invasive. Here, we showed the result for bladder cancer cells. The main findings indicate that cell mechanics is strongly related to the organization of the actin cytoskeleton, i.e., to the presence of thick actin bundles. The actin cytoskeleton is also related to migratory properties, indicating the presence of actin filaments, probably stress fibers. However, larger cell deformability is not related to the invasive properties of bladder cancer cells, which are shown here to be related to actin filaments. Our results demonstrate that the relation between migratory, invasive, and mechanical phenotypes has to be determined for each type of cancer cell. The potential applicability to identify cancerous cells by studying migratory and mechanical properties was demonstrated for the lectin-coated surfaces, which gives new pathways for surface modification for biosensors.

## Supplementary Materials

The following supporting information can be downloaded at www.mdpi.com/xxx/s1, Supplementary Notes S1, Supplementary Figure S1: *Transmigration of bladder cancer cells*; Supplementary Figure 2: *Proliferation of bladder cancer cells on lectin-coated surfaces*; Supplementary Table S1: *Migration speed determined for bladder cancer cells cultured on a lectin-coated surface*; Supplementary Table S2: *Young’s modulus was determined for bladder cancer cells cultured on a lectin-coated surface*.

## Author Contributions

Marcin Luty - Formal analysis, Methodology, Investigation, Validation, Writing – writing initial draft, review & editing

Renata Szydlak - Methodology, Investigation, Validation, Writing – review & editing

Joanna Pabijan - Methodology, Investigation, Validation, Writing – review & editing

Victorien E. Prot - Formal analysis, Methodology, Investigation, Validation, Writing – review & editing

Ingrid H. Øvreeide - Formal analysis, Methodology, Investigation, Validation, Writing – review & editing

Joanna Zemła - Formal analysis, Methodology, Investigation, Validation, Writing – review & editing

Bjørn T. Stokke - Supervision, Conceptualization, Validation, Funding acquisition, Writing – review & editing

Małgorzata Lekka - Supervision, Conceptualization, Resources, Funding acquisition, Writing – writing initial draft, Writing – review & editing

## Data Availability Statement

The data presented in this study are available on request from the corresponding author.

## Supplementary Figure S1: Migration trajectories of single cells

**Supplementary Figure S1.**
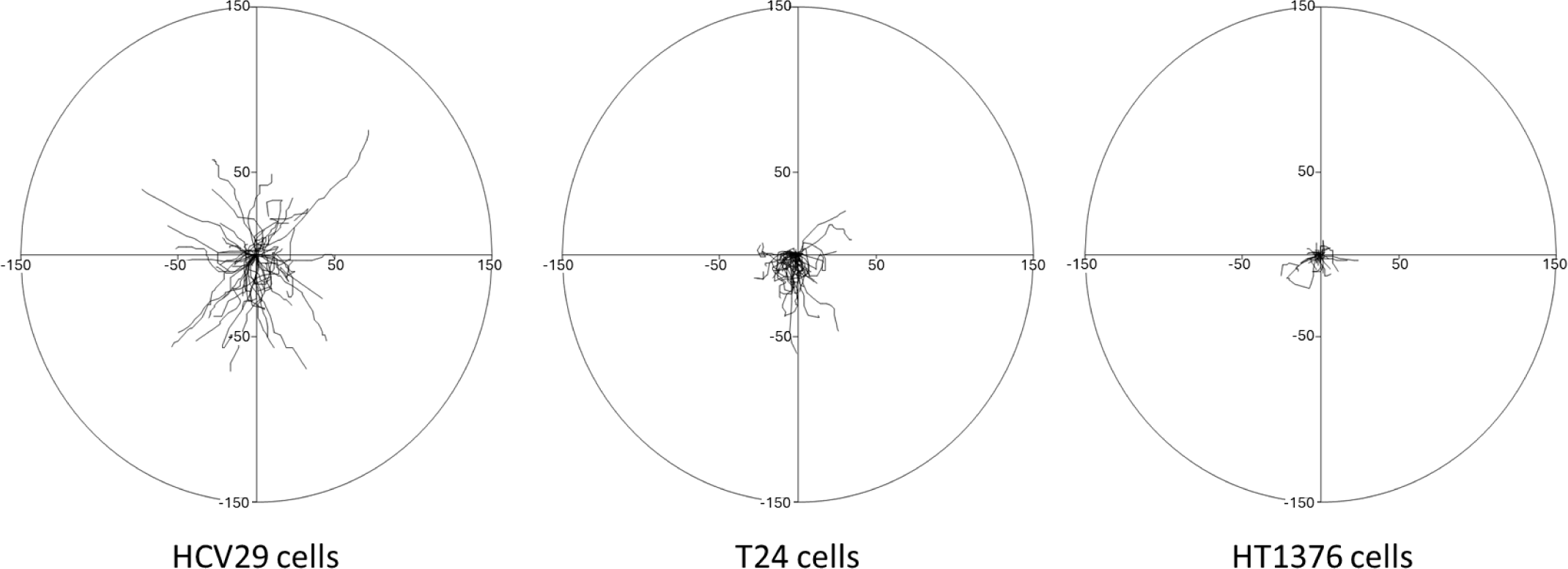
Migration trajectories of single bladder cancer cells. Exemplary trajectories were recorded for single non-malignant HCV29, invasive T24, and non-invasive HT1376 cells (X Y axes are in microns). The means displacements were 37.52 ± 1.94 µm (n = 150 cells), 16.38 ± 2.34 µm (n = 150 cells), and 7.33 ± 3.08 µm (n = 150 cells), respectively.

## Supplementary Note S1: Transmigration of cancer spheroids through HCV29 cell monolayer

The surface of the Corning Transwells inserts (Sigma-Aldrich, Poland, Poznań) with pore sizes of 8 µm was coated in two ways. First, a solution of ECM gel from Engelbreth-Holm-Swarm murine sarcoma (300 µg/mL, Sigma-Aldrich, Poznań, Poland; referred to here as matrigel) was placed (100 ul per traswell) and incubated at 37°C for 3 hours. The solution was removed, and the inserts were dried in a laminar flow chamber (Nuaire, USA). In separate experiments, the surface of the inserts was coated with a monolayer of non-malignant HCV29 cells. In this case, 100 µl of the cell solution (5×10^4^ cells were suspended in 100 µl culture medium supplemented with 10% serum) was added, followed by incubation at 37°C in the CO_2_ incubator for 24 hours. To form cancer spheroids, 4000 cells/well and 6000 cells/well for HT1376 and T24 cells were placed in non-adherent Nunclon Sphera 96-well plates. Then, after 96 hours, the formed spheroids were transferred to the Eppendorf tube (1.5 mL) using a Pauster pipette and incubated with 5 µM Cell Tracker Red CMPTX dye (Invitrogen, Thermo Fisher Scientific, Waltham, MA, USA) for 30 minutes. Afterward, cancer spheroids were washed with PBS buffer, suspended in the RPMI 1640 or EMEM medium containing 1% FBS, and placed in the upper Transwell chamber (coated with a monolayer of HCV29 cells). The lower Transwell chamber was filled with the RPMI 1640 or EMEM medium containing 10% FBS. Then, after 24 h, images of the lower chamber surface with cells were recorded using a fluorescence microscope.

**Supplementary Figure S2.**
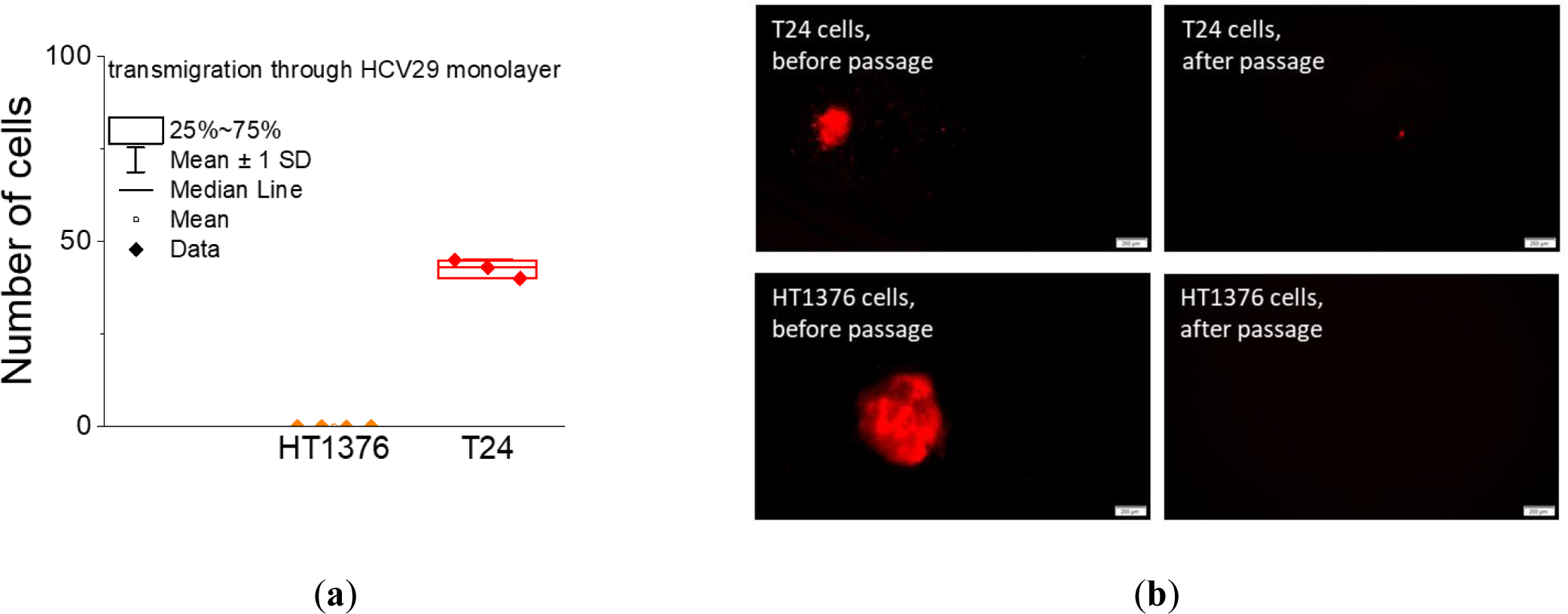
Transmigration of bladder cancer cells. **(a)** Transmigration of bladder cancer cells through the monolayer of non-malignant HCV29 cells. Each point denotes the total number of cells that migrated through one transwell, n = 3 repetitions. **(b)** Fluorescent images of spheroids formed from cancerous HT1376 and T24 cells, labeled with Cell Tracker Red dye, before and after transmigration through the monolayer of HCV29 cells. The scale bar is 200 µm.

**Supplementary Figure S3.**
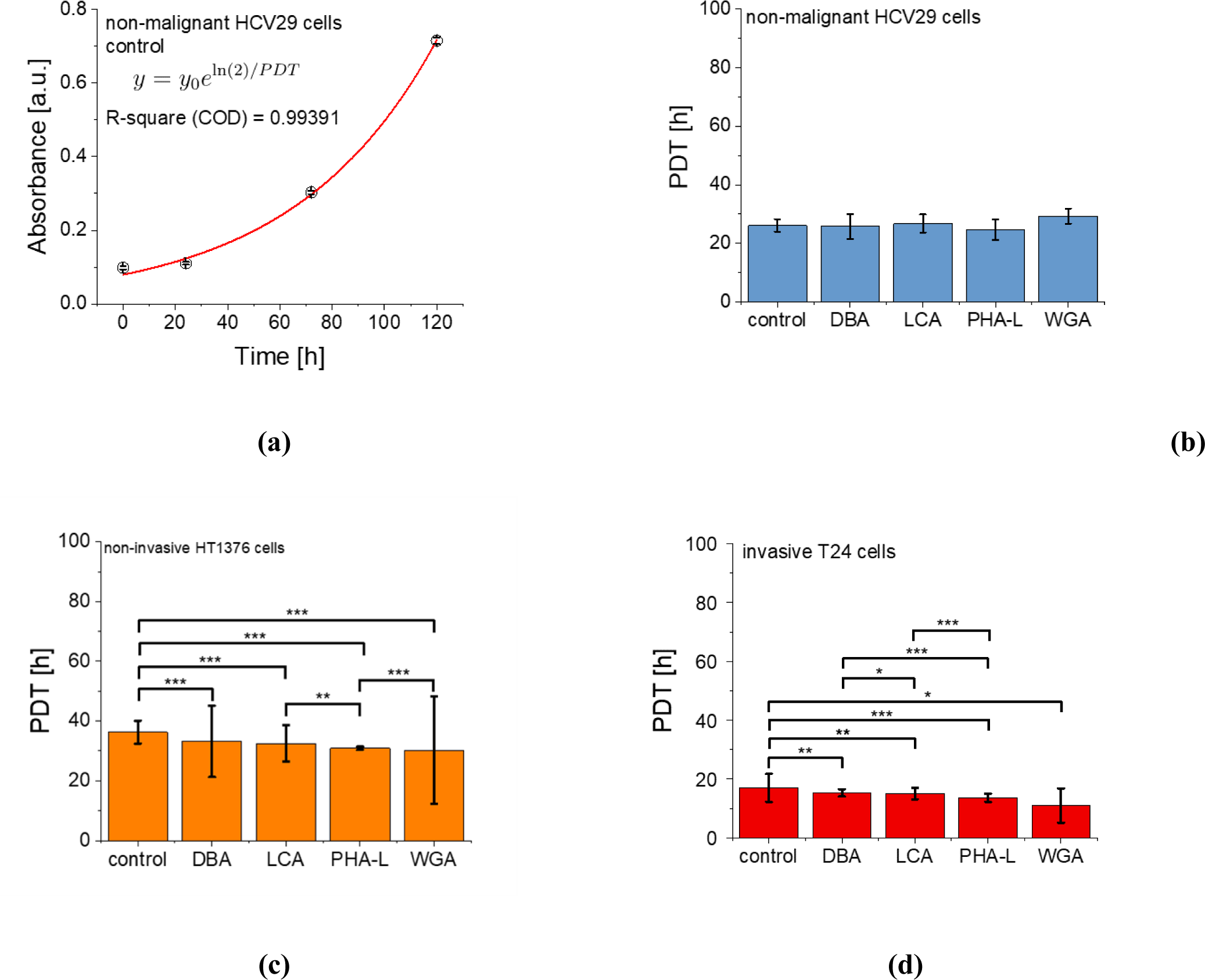
Proliferation of bladder cancer cells on lectin-coated surfaces. PDT time was obtained on lectin-coated (DBA, LCA, PHA-L, and WGA) surfaces for bladder cancer cells. (a) Exemplary data with the fit used for the PDT determination. (b) non-malignant HCV29, (c) non-invasive HT1376, and (d) invasive T24 cells. The bare surface of the plastic Petri dish was used as a control. Error bars are parameter standard errors originating from the fit. The F-test was used to determine whether the two datasets fitted with the same model differed significantly. Notation: ns – not statistically significantly different, *p < 0.05, **p < 0.01, ***p < 0.001.

**Supplementary Table S1.** Migration speed determined for bladder cancer cells cultured on a lectin-coated surface (expressed in um/min). The number of analyzed cells *n* = 150 cells (values statistically significant from the control are shown in red, based on a non-parametric Mann-Whitney test at a significance level of 0.05).

| Cells | control | DBA | LCA | PHA-L | WGA |
| --- | --- | --- | --- | --- | --- |
| non-malignant HCV29 | $0.24 \pm 0.11$ | $0.21 \pm 0.07$ | $0.20 \pm 0.08$ | $0.24 \pm 0.09$ | $0.18 \pm 0.10$ |
| invasive T24 | $0.15 \pm 0.11$ | $0.15 \pm 0.13$ | $0.14 \pm 0.12$ | $0.12 \pm 0.12$ | $0.07 \pm 0.08$ |
| non-invasive HT1376 | $0.05 \pm 0.06$ | $0.03 \pm 0.39$ | $0.03 \pm 0.38$ | $0.02 \pm 0.02$ | $0.03 \pm 0.04$ |

**Supplementary Table S2.**
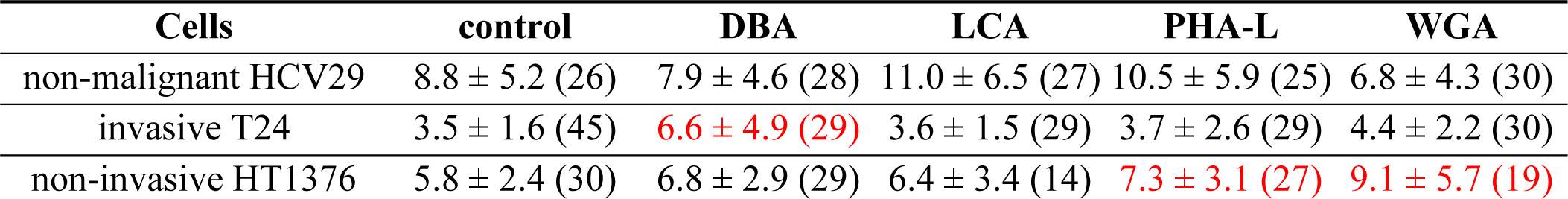
Young’s modulus was determined for bladder cancer cells cultured on a lectin-coated surface (expressed in kPa). Number of analyzed cells varied from 14 to 45 (shown in () in the table). Values statistically significant from the control are shown in red, based on the non-parametric Mann-Whitney test at a significance level of 0.05.

